# Extended Amygdala CRF Projections to Striatal Striosomes Potentiate Dopamine Suppression During Fentanyl Withdrawal

**DOI:** 10.64898/2026.06.22.733757

**Authors:** Xueyi Xie, Amanda Essoh, Ruifeng Chen, Xuehua Wang, Jun Wang

## Abstract

Opioid withdrawal is a powerful negative affective state that promotes continued drug use and relapse, and is often associated with hypodopaminergic states. Although corticotropin-releasing factor (CRF) signaling in the extended amygdala has been strongly implicated in stress and negative affect, how withdrawal-recruited CRF systems suppress dopamine release remains poorly understood. Here, we identify an extended amygdala–striosome–dopamine pathway engaged during fentanyl withdrawal. Naloxone-precipitated fentanyl withdrawal selectively recruited CRF neurons in the central amygdala and bed nucleus of the stria terminalis. These CRF neurons provided monosynaptic input to dorsostriatal medium spiny neurons (MSNs). Because striosomal MSNs are positioned to regulate dopamine release, we next found that CRF enhanced glutamatergic synaptic transmission onto striosomal MSNs, with CRF receptor 1 (CRFR1) signaling required for the postsynaptic strengthening of excitatory transmission. In vivo dopamine photometry further revealed that naloxone-precipitated withdrawal amplified striosome-mediated suppression of dopamine release, and this amplification was prevented by CRFR1 antagonism. Together, these findings reveal a mechanism by which withdrawal-recruited CRF signaling potentiates striosomal control of dopamine release. This extended amygdala–striosome–dopamine pathway may contribute to the hypodopaminergic negative affective state that drives opioid withdrawal and relapse vulnerability.

**Highlights:**

- Fentanyl withdrawal selectively recruits CeA and BNST CRF neurons
- CeA and BNST CRF neurons provide monosynaptic input to striatal MSNs
- CRF potentiates glutamatergic transmission onto striosomal MSNs via CRFR1
- CRFR1 signaling amplifies striosome-mediated dopamine suppression during withdrawal

## Introduction

Opioid use disorder is maintained not only by the positive reinforcing effects of opioids, but also by the powerful negative affective state that emerges during withdrawal [1–3]. In this negative reinforcement process, individuals continue or resume opioid use to relieve withdrawal-associated aversion, dysphoria, and motivational distress. A major candidate system for generating withdrawal-related negative affect is CRF signaling in the extended amygdala [4–7]. The central nucleus of the amygdala (CeA) and bed nucleus of the stria terminalis (BNST) are core components of this stress-related network and undergo marked neuroadaptations during opioid withdrawal and prolonged abstinence[6, 8]. Previous studies have shown that suppression of intra-amygdala CRF signaling weakens the aversive motivational effects of morphine withdrawal, and CRFR1 antagonism can block withdrawal-induced conditioned place aversion [9, 10]. These findings support the idea that extended amygdala CRF signaling is a key mediator of the aversive state of opioid withdrawal.

Opioid withdrawal has been associated with hypodopaminergic and amotivational states, which are thought to contribute to negative affect and relapse vulnerability [2, 3]. However, how withdrawal-recruited CRF signaling is translated into changes in dopamine-dependent motivational state remains unclear. The dorsal striatum is well-positioned to participate in this process because it integrates limbic, cortical, and neuromodulatory signals to regulate motivated behavior [11–13]. Within the dorsal striatum, the striosomal compartment is particularly relevant to opioid withdrawal and dopamine regulation. Striosomes are neurochemically specialized compartments enriched for μ-opioid receptor expression and preferentially associated with limbic-related inputs [12, 13]. They also provide a direct striatal route for GABAergic regulation of substantia nigra pars compacta (SNc) dopamine neurons, positioning them to convert affective and opioid-sensitive signals into changes in dopaminergic output [14–16]. Importantly, recent work showed that the activity of striosomal neurons causally controls fentanyl withdrawal-induced physical and anxiety-like symptoms, and that chronic opioid exposure can trigger GABAergic striatonigral plasticity capable of producing a hypodopaminergic state [17]. Together, these findings suggest that MOR-enriched striosomal circuits may be a critical substrate through which opioid withdrawal suppresses dopamine signaling.

A key unresolved question is what upstream withdrawal-related signal amplifies striosomal control of dopamine release. CRF signaling is a strong candidate because aversive and stress-related states robustly recruit it and has emerging roles in modulating dorsal striatal function[18–20]. Recent work identified CRF-positive inputs from the CeA and BNST to dorsal striatal cholinergic interneurons and showed that CRF enhances cholinergic interneuron excitability through CRFR1-dependent mechanisms [18, 19]. This suggests that extended amygdala CRF neurons may influence dorsal striatal function via multiple cellular targets. Yet whether withdrawal-recruited CRF systems directly modulate striatal projection neurons, especially striosomal MSNs, and whether this modulation alters dopamine release during opioid withdrawal remain unknown.

Here, we tested the hypothesis that fentanyl withdrawal recruits extended amygdala CRF signaling to amplify striosome-mediated suppression of dopamine release. We focused on CRF neurons in the CeA and BNST, their functional connectivity with dorsostriatal MSNs, the CRFR1-dependent modulation of excitatory transmission onto striosomal MSNs, and the impact of CRFR1 signaling on striosome-driven dopamine suppression during naloxone-precipitated fentanyl withdrawal. This study identifies an extended amygdala–striosome–dopamine mechanism through which CRF signaling may transform opioid withdrawal into a hypodopaminergic negative affective state.

## Results

### Naloxone-precipitated fentanyl withdrawal selectively recruits CRF ensembles in the extended amygdala

We first asked whether CRF systems within the extended amygdala are selectively engaged during fentanyl withdrawal. To address this, we generated CRF-Cre;Ai14 mice, in which CRF-expressing neurons are labeled with tdTomato (Fig. 1A). Mice underwent a 5-day fentanyl administration paradigm with escalating doses (Fig. 1B). Locomotor activity was monitored using infrared beam breaks within a plexiglass chamber.

**Figure 1.**
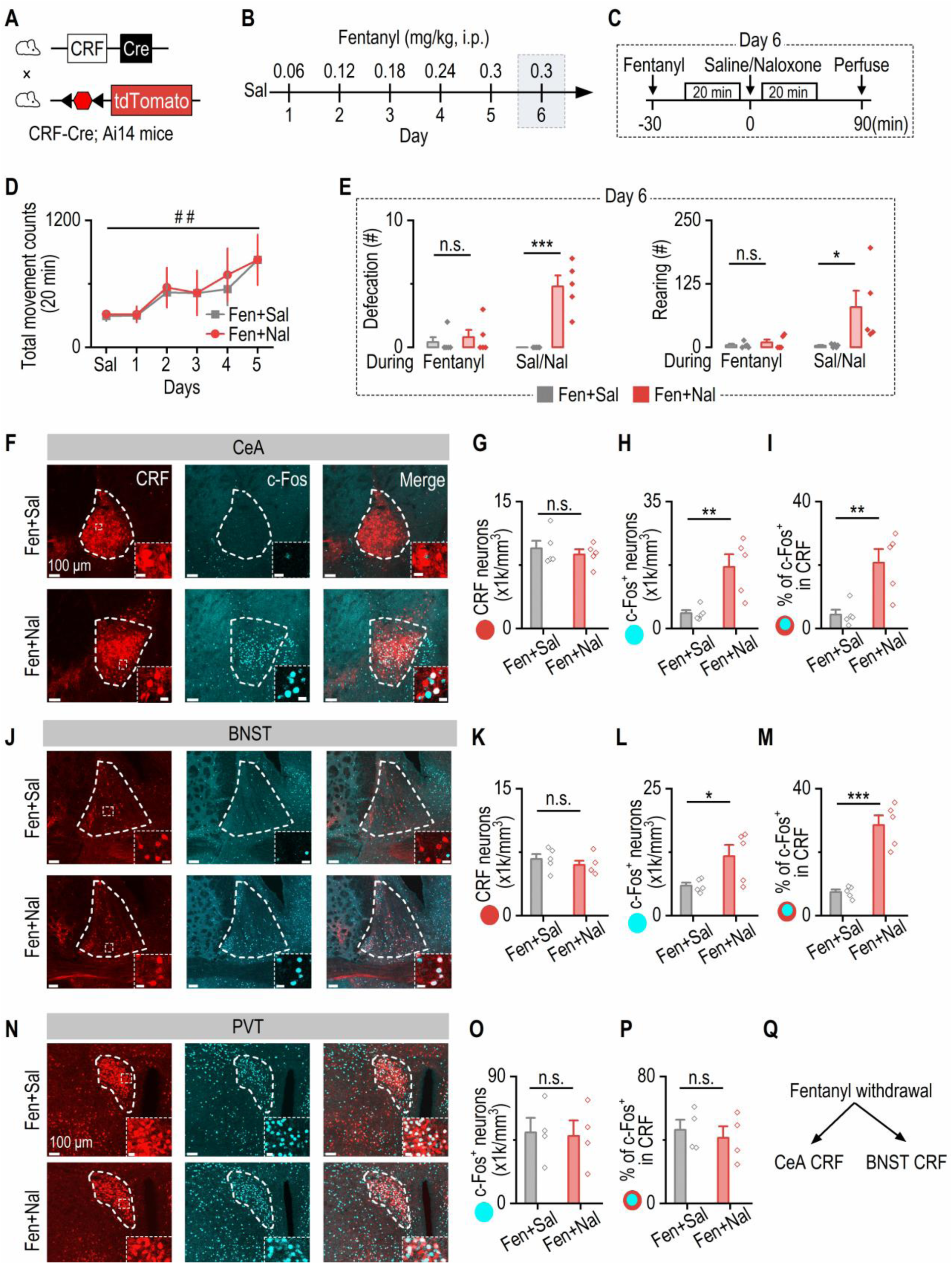
Fentanyl withdrawal recruits CRF ensembles in the CeA and BNST, but not PVT. **A**, Breeding strategy used to generate CRF-Cre;Ai14 mice, in which CRF+ neurons express tdTomato. **B**, Experimental timeline. Mice were first habituated to saline injections, then received five days of escalating fentanyl injections. **C**, Day 6 withdrawal paradigm. Ten minutes after fentanyl injection, mice were placed in an open-field arena for 20 min. They then received either saline or naloxone and were monitored for an additional 20 min. Ninety minutes after the saline/naloxone injection, brains were collected for c-Fos immunohistochemistry. **D**, Total locomotor activity increased progressively across the 5-day fentanyl regimen. Two-way RM ANOVA: F_(4,32)_ = 4.59, ^##^p = 0.0049. **E**, Behavioral signs of precipitated withdrawal. Left: Defecation counts were similar between groups during fentanyl exposure but increased selectively following naloxone (Two-way RM ANOVA: Group: F_(1,8)_ = 64.38, p < 0.001. Sidak’s post hoc test: Fen+Sal vs. Fen+Nal during Fen: t = 0.51, p = 0.85; Fen+Sal vs. Fen+Nal during Sal/Nal: t = 6.096,***p < 0.001). Right: Rearing behavior showed the same pattern, increasing only in naloxone-treated mice (Two-way RM ANOVA: Group: F_(1,8)_ = 7.29, p = 0.027. Sidak’s post hoc test: Fen+Sal vs. Fen+Nal during Fen: t = 0.23, p = 0.97; Fen+Sal vs. Fen+Nal during Sal/Nal: t = 3.18, *p = 0.011). **F**, Representative confocal images of CRF+ (red) and c-Fos+ (cyan) neurons in the CeA after saline or naloxone administration. Insets show higher magnification of boxed regions. Scale bar for the inset: 20 μm. **G-I**, Quantification in the CeA. (G) CRF+ neuron density did not differ between groups (unpaired t test: t_(8)_ = 0.65, p = 0.53). (H) Naloxone significantly increased the density of c-Fos+ neurons (unpaired t test: t_(8)_ = 3.58, **p = 0.0072). (I) A greater proportion of CRF+ neurons were c-Fos+ in naloxone-treated mice (unpaired t test: t_(8)_ = 3.59, **p = 0.0071). **J**, Representative confocal images of CRF+ and c-Fos+ neurons in the BNST. Scale bar for the inset: 20 μm. **K-M**, Quantification in the BNST. (K) CRF+ neuron density was similar between groups (unpaired t test: t_(8)_ = 0.88, p = 0.40). (L) Naloxone increased c-Fos+ neuron density (unpaired t test: t_(8)_ = 2.51, *p = 0.036). (M) The percentage of CRF+ neurons expressing c-Fos was markedly elevated in naloxone-treated mice (unpaired t test: t_(8)_ = 6.69, ***p = 0.0002). **N**, Representative confocal images of CRF+ and c-Fos+ neurons in the PVT. Scale bar for the inset: 20 μm. **O**, **P**, Quantification in the PVT. (O) Naloxone did not alter c-Fos+ neuron density (unpaired t test: t_(6)_ = 0.16, p = 0.88). (P) The percentage of CRF+ neurons expressing c-Fos was similar between groups (unpaired t test: t_(6)_ = 0.52, p = 0.62). **Q**, Summary schematic. n = 5 mice per group (D-M) and 4 mice per group (O-P).

On day 6, mice received fentanyl and were monitored for 20 minutes, followed by injection of either saline or naloxone to induce precipitated withdrawal (Fig. 1C). Behavioral responses were recorded for an additional 20 minutes, after which animals were perfused 90 minutes later for c-Fos immunohistochemistry to identify neurons activated during withdrawal.

Repeated fentanyl exposure progressively increased locomotor activity (Fig. 1D), indicating effective systemic drug action. To assess withdrawal-related behaviors, we quantified defecation and rearing, which reflect somatic withdrawal symptoms and negative affect-like states, respectively. During fentanyl exposure, mice exhibited minimal defecation and rearing (Fig. 1E). In contrast, naloxone administration robustly increased both defecation and rearing behavior compared to saline controls. These results confirm the induction of pronounced physical and affective withdrawal symptoms by naloxone.

We next examined neuronal activation within major CRF-rich regions, including the central amygdala (CeA), bed nucleus of the stria terminalis (BNST), and paraventricular thalamus (PVT), using c-Fos immunolabeling. In the CeA, naloxone did not alter the density of CRF-expressing neurons (Fig. 1F, 1G), but significantly increased the number of c-Fos⁺ neurons (Fig. 1H), resulting in a higher proportion of CRF neurons being activated (Fig. 1I). A similar pattern was observed in the BNST: CRF neuron density remained unchanged (Fig. 1J, 1K), whereas overall neuronal activation was significantly elevated following naloxone administration (Fig. 1L), leading to increased recruitment of CRF neurons (Fig. 1M). In contrast, the PVT showed no significant changes. Neither overall neuronal activation (Fig. 1N, 1O), nor the proportion of CRF neurons activated (Fig. 1P) differed between saline and naloxone groups.

Together, these findings demonstrate that naloxone-precipitated fentanyl withdrawal selectively recruits CRF neuronal ensembles in the extended amygdala—specifically within the CeA and BNST—but not in the PVT (Fig. 1Q).

### Withdrawal-relevant CRF neurons provide monosynaptic input to CRFR1-expressing striatal MSNs

Our previous work has identified the dorsal striatum as a key locus mediating fentanyl withdrawal-related behaviors [17]. Having established that naloxone-precipitated withdrawal selectively recruits CRF ensembles in the CeA and BNST, we next asked whether these withdrawal-relevant CRF neurons directly project to dorsostriatal medium spiny neurons (MSNs).

To test this, we injected a Cre-dependent ChRmine virus into either the CeA or BNST of CRF-Cre mice, enabling selective expression of ChRmine in CRF neurons (Fig. 2A). We then performed whole-cell patch-clamp recordings from MSNs in the dorsal striatum while optically stimulating CRF axon terminals with 590-nm light. Photostimulation of CeA-derived CRF terminals evoked robust postsynaptic currents in striatal MSNs (Fig. 2B). These responses were abolished by bath application of tetrodotoxin (TTX, 1 μM), indicating dependence on action potential propagation, and were subsequently restored by co-application of TTX and 4-aminopyridine (4-AP, 500 μM), consistent with monosynaptic transmission. Similar results were observed for BNST-derived CRF projections (Fig. 2C), demonstrating that CRF neurons from both regions establish direct monosynaptic connections with dorsostriatal MSNs.

**Figure 2.**
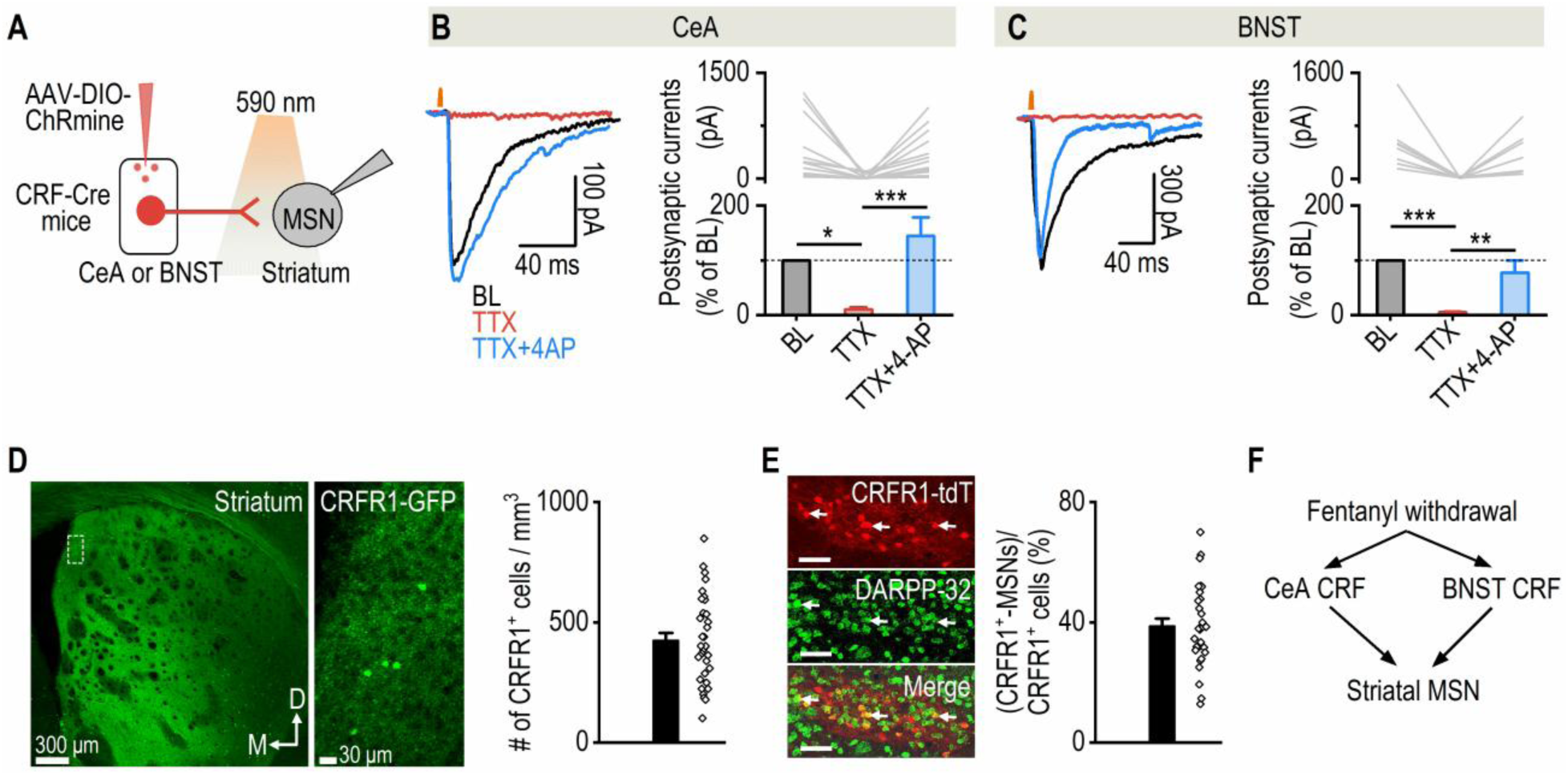
CeA and BNST CRF neurons monosynaptically target CRFR1-expressing striatal medium spiny neurons. **A**, Experimental schematic. CRF-Cre mice received bilateral infusions of AAV-DIO-ChRmine into either the CeA or BNST. Six weeks later, whole-cell patch-clamp recordings were performed in dorsal striatal MSNs. Synaptic transmission from CeA or BNST CRF⁺ projections to striatal MSNs was assessed by stimulating ChRmine-expressing axon terminals with 590-nm light. **B**, Optically evoked postsynaptic currents in striatal MSNs following stimulation of CeA CRF⁺ terminals were abolished by bath application of tetrodotoxin (TTX) and restored upon subsequent application of 4-aminopyridine (4-AP). One-way RM ANOVA: F_(2,24)_ = 11.69, p = 0.0003; Tukey’s multiple comparsions test: BL vs. TTX: q = 4.489, *p = 0.011; TTX vs TTX+4-AP: q = 6.712, ***p = 0.0002. n = 13 neurons / 3 mice. **C**, Optically evoked postsynaptic currents in striatal MSNs following stimulation of BNST CRF⁺ terminals were abolished by bath application of TTX and restored upon subsequent application of 4-AP. One-way RM ANOVA: F_(2,14)_ = 15.29, p = 0.0003; Tukey’s multiple comparsions test: BL vs. TTX: q = 7.481, ***p = 0.0003; TTX vs TTX+4-AP: q = 5.716, **p = 0.0033. n = 8 neurons / 2 mice. **D**, Representative confocal images and quantification showing CRFR1-expressing neurons in the dorsal striatum. n = 35 slices / 4 mice. **E**, Approximately 40% of striatal CRFR1-expressing neurons co-express DARPP-32. n = 30 slices / 4 mice. **F**, Summary schematic.

We next asked whether these postsynaptic targets express CRF receptors. Using CRFR1-GFP reporter mice [21, 22], we found that CRFR1-expressing neurons are distributed throughout the dorsal striatum, albeit sparsely (Fig. 2D). To determine their cellular identity, we utilized CRFR1-tdTomato mice and co-labeled sections with DARPP-32, a marker for MSNs. Approximately 40% of CRFR1-expressing neurons co-expressed DARPP-32, indicating that a substantial fraction of CRFR1⁺ cells are MSNs (Fig. 2E).

Together, these results demonstrate that CRF neurons in the CeA and BNST form monosynaptic projections onto dorsostriatal MSNs (Fig. 2F), many of which express CRFR1, providing a potential anatomical and molecular substrate for CRF modulation of striatal function during withdrawal.

### CRFR1 signaling potentiates excitatory transmission in striosomal MSNs

Having shown that withdrawal-recruited CRF neurons from the CeA and BNST provide direct input to striatal MSNs, we next asked whether CRF signaling directly modulates striosomal MSN activity. This question is particularly relevant because our previous work identified striosomal neurons as key mediators of aversion, negative reinforcement, and fentanyl withdrawal-related symptoms[17, 23, 24]. Since MSNs are largely quiescent at baseline and depend on glutamatergic inputs to drive neuronal activity, we examined whether CRF alters excitatory synaptic transmission onto striosomal neurons.

To selectively target striosomal neurons, we generated MOR-Cre;Ai14 mice, in which MOR-expressing striosomal neurons are labeled with tdTomato (Fig. 3A). We then performed whole-cell patch-clamp recordings from tdTomato⁺ striosomal MSNs and measured spontaneous excitatory postsynaptic currents (EPSCs) under different pharmacological conditions (Fig. 3B).

**Figure 3.**
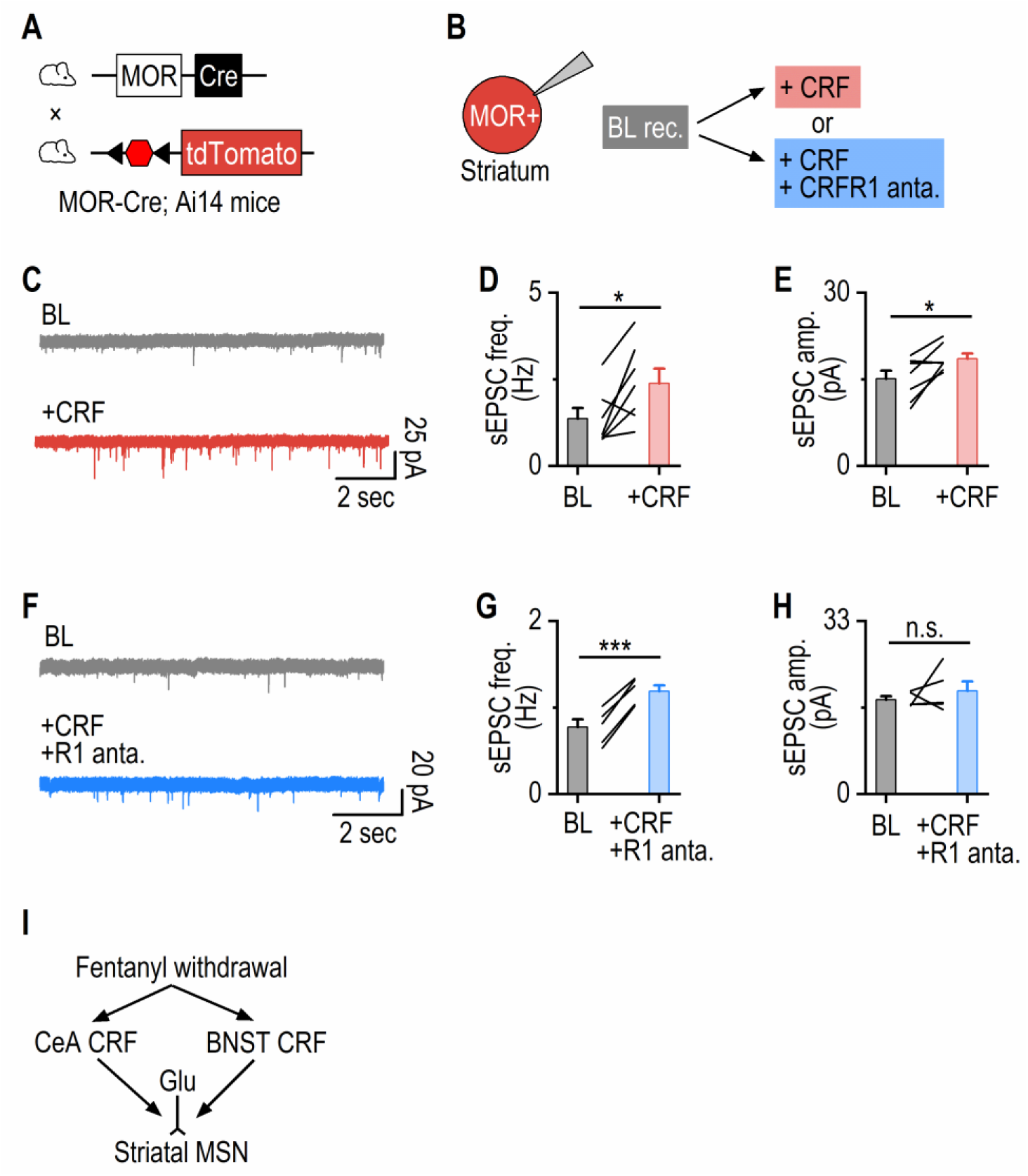
CRFR1 signaling enhances the excitatory input of striatal striosomal MSNs. **A**, Schematic illustrating the breeding strategy of MOR-Cre;Ai14 mouse to visualize striatal striosomal neurons. **B**, Schematic illustrating the recording procedure. Whole-cell patch clamp recordings were conducted in tdT+ striosomal neurons. Once neurons were patched, a 3-6 min baseline (BL) spontaneous excitatory poststynaptic currents (sEPSCs) were recorded. After that, either CRF or a CRF+CRFR1 antagonist was bath-applied, and another 6 min of sEPSCs were recorded. **C**, Sample traces of sEPSCs during baseline and CRF bath application. **D**, **E**, CRF bath application (100 nM) increased both the frequencies (D; paired t test: t_(6)_ = 2.85, *p = 0.029) and amplitudes (E; paired t test: t_(6)_ = 3.14, *p = 0.020) of sEPSCs in striosomal neurons. n = 7 neurons / 3 mice. **F**, Sample traces of sEPSCs during baseline and CRF+CRFR1 antagonist bath application. **G**, **H**, CRF+CRFR1 antagonist bath application increased the frequencies (G; paired t test: t_(4)_ = 11.53, ***p = 0.0003), but not amplitudes (H; paired t test: t_(4)_ = 0.81, p = 0.46), of sEPSCs in striosomal neurons. n = 5 neurons / 3 mice. **I**, Summary schematic

Bath application of CRF significantly increased both the frequency and amplitude of spontaneous EPSCs in striosomal MSNs (Fig. 3C-3E). The increase in sEPSC frequency suggests enhanced presynaptic glutamate release onto striosomal neurons, whereas the increase in sEPSC amplitude indicates potentiation of postsynaptic glutamatergic responsiveness. We next asked whether these CRF-induced effects were mediated by CRFR1 signaling. To test this, we co-applied CRF with a CRFR1 antagonist. Under CRFR1 blockade, CRF still increased sEPSC frequency (Fig. 3F, 3G), but no longer increased sEPSC amplitude (Fig. 3H). These results suggest that the CRF-induced enhancement of presynaptic glutamate release may involve CRFR1-independent mechanisms, whereas the postsynaptic potentiation of excitatory transmission is dependent on CRFR1 signaling.

Together, these findings demonstrate that CRF enhances glutamatergic transmission onto striosomal MSNs through both presynaptic and postsynaptic mechanisms. Importantly, CRFR1 signaling is required for the postsynaptic strengthening of excitatory transmission in striosomal MSNs, providing a potential mechanism by which withdrawal-recruited CRF inputs may amplify striosomal activity during fentanyl withdrawal (Fig. 3I).

### CRFR1 signaling amplifies striosomeal suppression of dopamine release during fentanyl withdrawal

Our previous work showed that fentanyl withdrawal potentiates striosome-mediated inhibition of dopamine release, and that this enhanced striosome–dopamine inhibition may contribute to withdrawal symptoms[17]. In the present study, we found that naloxone-precipitated fentanyl withdrawal recruits CRF neurons in the CeA and BNST, that these CRF neurons provide monosynaptic input to striatal MSNs, and that CRF potentiates excitatory transmission onto striosomal MSNs. Together, these findings led us to hypothesize that withdrawal-recruited CRF signaling may be a mechanism that amplifies striosome-mediated suppression of dopamine release during fentanyl withdrawal.

To test this hypothesis in vivo, we used optogenetic activation of striosomal neurons as a controlled output assay. This approach was important because striatal MSNs, including striosomal MSNs, are generally quiescent under baseline conditions due to their hyperpolarized resting membrane potentials. Therefore, rather than relying on spontaneous striosomal activity, we activated striosomal neurons with the same light intensity across baseline, fentanyl, and naloxone-precipitated withdrawal conditions. This allowed us to determine whether the withdrawal state increases the functional ability of striosomal neurons to suppress dopamine release, and whether this potentiation depends on CRFR1 signaling.

To do this, we injected a Cre-dependent ChRmine into the DMS of MOR-Cre mice to selectively express ChRmine in striosomal neurons. In the same region, we co-expressed the dopamine sensor GRAB_DA3h_ to monitor dopamine dynamics in vivo (Fig. ^4^A, 4B). Optical fibers were implanted above the injection site, allowing simultaneous 590-nm stimulation of ChRmine-expressing striosomal neurons and fiber photometry recording of dopamine signals. Mice were tested sequentially under three conditions: baseline, fentanyl exposure, and naloxone-precipitated withdrawal (Fig. 4C). Before recording, mice received either saline or a CRFR1 antagonist to determine whether CRFR1 signaling contributes to withdrawal-induced potentiation of striosome-mediated dopamine suppression.

**Figure 4.**
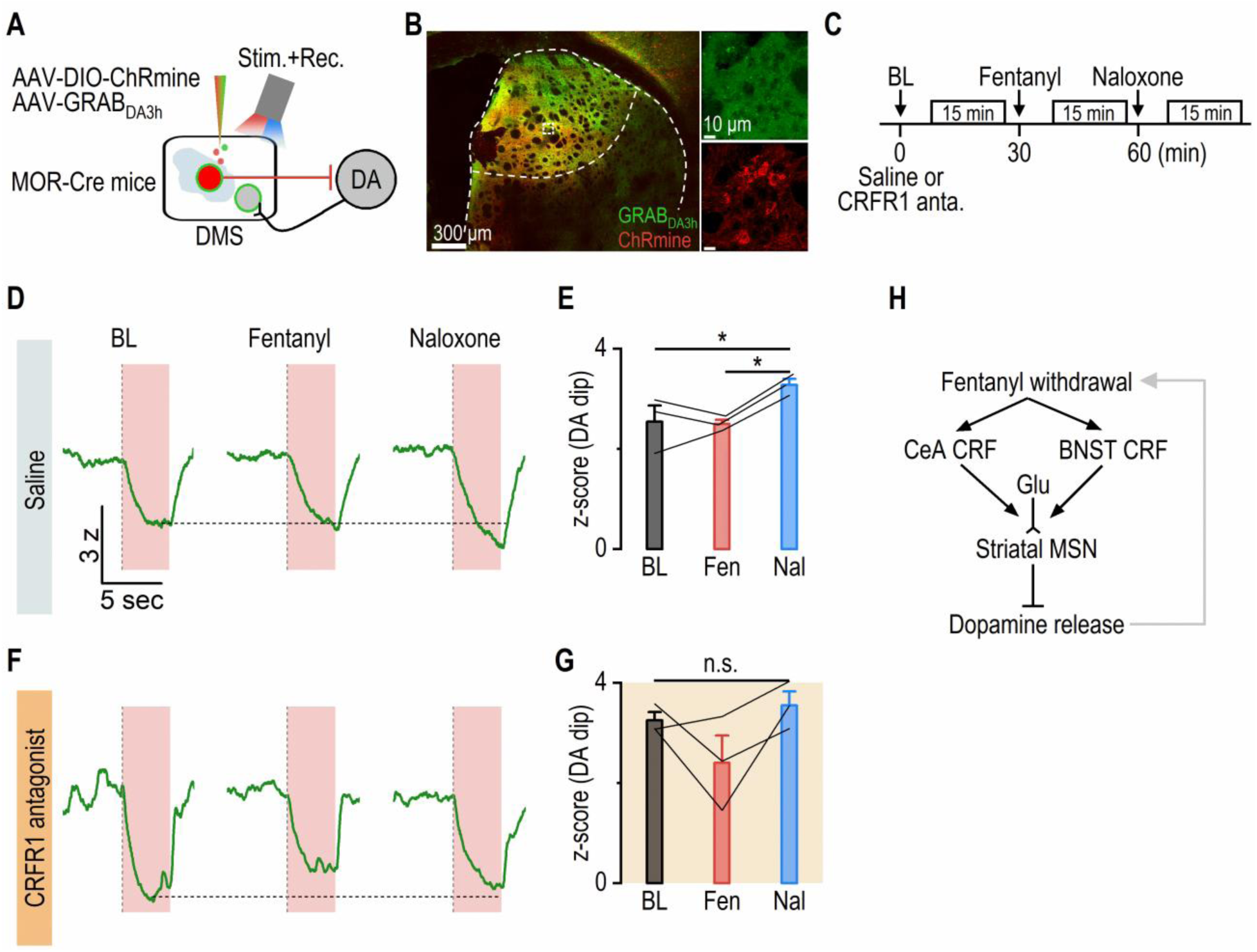
Fentanyl withdrawal amplifies striosome-mediated suppression of dopamine release, which is mediated by CRFR1 signaling. **A**, Experimental schematic. MOR-Cre mice received bilateral injections of AAV-DIO-ChRmine and AAV-GRAB_DA3h_ into the DMS. Optical fibers were implanted above the injection site for simultaneous optogenetic stimulation and dopamine recording. Red light (590 nm) was delivered to activate ChRmine-expressing MOR⁺ striosomal neurons, which inhibit midbrain dopaminergic neurons, while GRAB_DA3h_ fluorescence was used to monitor striatal dopamine dynamics. **B**, Sample confocal images of ChRmine and GRAB_DA3h_ expression in the DMS. **C**, Experimental timeline. Mice received saline or a CRFR1 antagonist injection. Ten minutes later, dopamine responses to MOR⁺ neuron stimulation were recorded for 15 min (baseline, BL). Fentanyl was then administered with dopamine dynamics recorded for 15 min, followed 30 min later by naloxone to precipitate withdrawal. Ten minutes after the naloxone injection, dopamine responses were recorded again for 15 min. **D**, **E**, Stimulation of MOR⁺ neurons produced a strong suppression of striatal dopamine release. This inhibitory effect was significantly greater during naloxone-precipitated withdrawal than at baseline. One-way RM ANOVA: F_(2,4)_ = 10.46, p = 0.0258. Tukey’s multiple comparisons test: BL vs Nal: q = 5.426, *p = 0.0396; Fen vs Nal: q = 5.763, *p = 0.0326. n = 3 mice. **F, G**, In mice pretreated with a CRFR1 antagonist, naloxone-precipitated withdrawal no longer enhanced dopamine suppression. Dopamine inhibition during withdrawal was comparable to baseline levels. One-way RM ANOVA: F_(2,4)_ = 2.931, p = 0.165. Tukey’s multiple comparisons test: BL vs Nal: q = 0.874, p = 0.8189; Fen vs Nal: q = 3.304, p = 0.161. n = 3 mice. **H**, Summary model. Fentanyl withdrawal activates CRF systems in both the CeA and BNST, which project to striatal MSNs, including striosomal populations. Enhanced activation of striosomal MOR⁺ MSNs increases inhibition of dopaminergic neurons, leading to amplified suppression of dopamine release during withdrawal. This effect is mediated by CRFR1 signaling.

In saline-treated mice, optogenetic activation of striosomal neurons induced a robust decrease in dopamine signal during the stimulation window (Fig. 4D). Acute fentanyl did not significantly alter the magnitude of this dopamine suppression. However, after naloxone administration, the same optogenetic stimulation produced a significantly larger dopamine dip (Fig. 4E), confirming that fentanyl withdrawal potentiates striosome-mediated inhibition of dopamine release.

We next asked whether this withdrawal-induced potentiation requires CRFR1 signaling. In mice treated with a CRFR1 antagonist, optogenetic activation of striosomal neurons still produced dopamine suppression (Fig. 4F), indicating that striosomal activation remained effective. However, naloxone no longer enhanced the magnitude of the dopamine dip (Fig. 4G). Thus, CRFR1 blockade prevented the withdrawal-induced potentiation of striosome-mediated dopamine suppression.

Together, these findings demonstrate that CRFR1 signaling is required for the enhancement of striosome-mediated dopamine inhibition during fentanyl withdrawal. Combined with our anatomical and electrophysiological findings, these results support a model in which withdrawal-recruited CeA and BNST CRF inputs potentiate striosomal MSN function through CRFR1 signaling, thereby amplifying striosome-mediated suppression of dopamine release during withdrawal (Fig. 4H).

## Discussion

Opioid withdrawal is not simply the absence of opioid reward; it is an actively generated negative affective state that promotes continued drug use and relapse. A major framework in addiction biology proposes that withdrawal recruits stress systems in the extended amygdala, including CRF signaling, which contributes to negative reinforcement and relapse vulnerability[25–28]. Our study extends this model by identifying a downstream striatal mechanism through which withdrawal-recruited CRF signaling may suppress dopamine release. Rather than acting only within classical stress circuits, CeA and BNST CRF neurons appear to engage a compartment-specific striatal output pathway, thereby linking extended amygdala stress signaling to striosome-mediated dopamine inhibition during fentanyl withdrawal. CRF systems have been strongly implicated in the withdrawal/negative affect stage of addiction[29], and opioid withdrawal is broadly understood to involve negative affective states that promote drug seeking and relapse[30].

A key implication of these findings is that the dorsal striatum may serve as an important interface between stress neuropeptide systems and dopamine-dependent motivational state. The CeA and BNST are well known for their roles in stress, anxiety, and aversive motivational processing, but their influence on dorsal striatal output has been less clear. Recent work showed that CRF-positive neurons in the CeA and BNST provide direct input to dorsal striatal cholinergic interneurons and modulate acetylcholine release through CRFR1 signaling [19]. Our study adds a complementary mechanism by showing that withdrawal-relevant CRF signaling can also influence striatal projection neurons, particularly striosomal MSNs. This expands the functional role of extended amygdala CRF projections from local stress regulation to direct modulation of striatal output circuits.

The striosome compartment provides a particularly compelling substrate for this effect. Striosomes are neurochemically distinct striatal compartments enriched for markers such as mu-opioid receptors and are positioned to regulate dopamine-containing neurons in the substantia nigra[13, 14, 31, 32]. Recent circuit work further supports the idea that striosomal pathways can directly modulate dopamine neurons and dopamine release, with striosomal D1 pathways exerting net inhibitory effects on dopamine-containing neurons[17, 33, 34]. Thus, the present findings suggest that withdrawal may recruit CRF signaling to increase the gain of a pre-existing striosome–dopamine “brake.” In this model, CRF does not create a new pathway during withdrawal; instead, it strengthens the functional impact of striosomal output on dopamine release.

Mechanistically, one important interpretation is that CRFR1 signaling may transform striosomal MSNs into a more excitable or more synaptically responsive state during withdrawal. Because MSNs are normally hyperpolarized and require coordinated excitatory input to fire, even modest enhancement of glutamatergic synaptic strength could substantially increase striosomal output. The CRF-induced increase in sEPSC amplitude suggests postsynaptic strengthening, while the persistence of the frequency effect under CRFR1 blockade suggests that presynaptic CRFR2 receptors or network-level mechanisms may also contribute[35]. A plausible postsynaptic mechanism is that CRFR1 activates cAMP/PKA- and/or ERK-dependent signaling, leading to enhanced AMPAR function, phosphorylation, or synaptic trafficking[35, 36].

Another important possibility is that CRF signaling may interact with opioid-induced plasticity in MOR-enriched striosomal neurons. Previous work showed that striatal MOR-positive neurons are strongly involved in fentanyl withdrawal-related physical and anxiety-like behaviors, and that repeated opioid exposure can alter direct-pathway output to dopamine-related circuits[17]. This suggests a two-hit model: chronic fentanyl exposure may prime MOR-enriched striosomal circuits, while naloxone-precipitated withdrawal recruits extended amygdala CRF signaling to further amplify striosomal output. Under this model, opioid withdrawal symptoms emerge not only from reduced opioid receptor activation, but from an active convergence of opioid-adapted striosomal circuits and stress-driven CRF modulation.

The selectivity of CRF recruitment is also conceptually important. The lack of strong PVT CRF recruitment suggests that fentanyl withdrawal does not simply activate all CRF-rich regions uniformly. Instead, withdrawal appears to preferentially engage CeA and BNST CRF ensembles, consistent with the central role of the extended amygdala in negative affective states. This regional selectivity supports a more precise model in which different CRF systems may contribute to different components of withdrawal. CeA/BNST CRF projections may be especially important for coupling aversive internal states to striatal dopamine suppression, whereas other CRF populations may regulate distinct physiological, cognitive, or motivational aspects of withdrawal.

The in vivo dopamine data further suggest that CRFR1 acts as a state-dependent gain mechanism rather than a basic requirement for striosome–dopamine connectivity. CRFR1 antagonism did not eliminate dopamine suppression produced by striosomal activation, but prevented the withdrawal-induced enhancement of this suppression. This distinction is important therapeutically. It suggests that CRFR1 blockade may selectively reduce the pathological amplification of dopamine suppression during withdrawal while leaving baseline striosome–dopamine communication relatively intact. This may help explain why targeting stress-related CRF signaling could be useful for withdrawal-related negative affect, even if CRFR1 signaling is not the sole determinant of dopamine tone.

Overall, our study provides a mechanistic bridge between three previously related but often separately studied systems: extended amygdala CRF signaling, striosomal MSN function, and dopamine suppression during opioid withdrawal. We demonstrate that withdrawal-recruited CRF signaling can be routed through a defined striosomal output pathway to amplify inhibition of dopamine release. This supports a model in which fentanyl withdrawal engages CeA/BNST CRF neurons to potentiate excitatory drive onto striosomal MSNs through CRFR1-dependent postsynaptic mechanisms, thereby enhancing striosome-mediated dopamine suppression. By linking stress neuropeptide signaling to compartment-specific striatal control of dopamine, this work provides a circuit mechanism through which opioid withdrawal may generate negative affective states that promote relapse.

## Materials and Methods

### Reagents

AAV5-Ef1a-DIO-hsChRmine-oScarlet was purchased from the UNC Vector Core, and AAV-GRAB_DA3h_ was purchased from BrainVTA. CRF peptide (#1151) and CRFR1 antagonist (NBI 35695; #3100) were purchased from Tocris. DARPP-32 (PA5-85788) antibodies were purchased from Thermo Fisher Scientific.

### Animals

Male and female 4- to 6-month-old mice were used in all studies. CRF-ires-CRE (stock 012704), Ai14 (stock 007914) and C57BL/6J (stock 000664) mice were purchased from The Jackson Laboratory. CRFR1-GFP mice were gifted by Dr. Marisa Roberto’s lab [21, 22]. CRFR1-tdTomato mice were gifted by Dr. Nicholas Justice’s lab[37]. MOR-Cre mice were gifted by Dr. Brigitte Kieffer’s lab [38]. CRF-Cre;Ai14 mice were generated by crossing CRF-Cre with Ai14 mice and MOR-Cre;Ai14 mice were generated by crossing MOR-Cre with Ai14 mice. Genotypes were confirmed through PCR analysis of tail DNA to detect Cre or fluorescent protein genes in mice. Animals were housed in a temperature-and humidity-controlled vivarium with a 12-h light/dark cycle. Food and water were available *ad libitum*. The Texas A&M University Institutional Animal Care and Use Committee approved all animal care and experimental procedures.

### Stereotaxic virus infusion

The stereotaxic virus infusion procedure was conducted as described previously [17, 19, 39–47]. AAV-DIO-ChRmine was infused into the CeA (AP: −1.60 mm; ML: ±4.20 mm; DV: −8.10 mm) or BNST (AP: −0.10 mm; ML: ±1.40 mm; DV: −6.70 mm) of CRF-Cre mice. AAV-DIO-ChRmine and AAV-GRAB_DA3h_ were co-infused into the DMS (AP: 0.3 mm, ML: ±1.65 mm, DV: −2.95 mm) of MOR-Cre mice. The animals were placed on a stereotaxic surgical frame after being sedated with 3-4% isoflurane at a rate of 1.0 L/min, as described previously. These coordinates were obtained from previous publications and verified using the mouse brain atlases. A volume of 0.5 µL/site was infused at a rate of 0.12 µL/min. At the end of the infusion, the injectors remained at the injection site for 10 to 15 min before removal to allow for virus diffusion. For MOR-Cre mice, optical fibers were implanted along with the injection track after a wait and at the DV of −2.90 mm. The scalp incision was then sutured, and the animals were returned to their home cage for recovery.

### Fentanyl administration and naloxone-precipitated withdrawal

Mice were habituated to intraperitoneal saline injections for two days before the fentanyl treatment paradigm. To induce fentanyl dependence, mice received once-daily intraperitoneal injections of escalating doses of fentanyl over 5 days: 0.06, 0.12, 0.18, 0.24, and 0.30 mg/kg on days 1–5, respectively. Ten minutes after each fentanyl injection, mice were placed in a neutral behavioral chamber equipped with a one-axis infrared beam-break system positioned on the floor. Locomotor activity was recorded as the number of beam breaks during the monitoring period.

On day 6, mice received 0.30 mg/kg fentanyl and were placed in the neutral behavioral chamber. Behavior was video recorded for 20 min before withdrawal induction. Mice then received an intraperitoneal injection of either saline or naloxone (5 mg/kg) to precipitate withdrawal, and behavior was recorded for an additional 20 min. Animals used for c-Fos analysis were perfused 90 min after saline or naloxone injection.

Withdrawal-related behaviors were quantified from video recordings. Defecation was assessed by counting the number of fecal boli after the 20-min post-fentanyl, post-saline, or post-naloxone monitoring period. Rearing behavior was manually scored from video recordings by an undergraduate experimenter blinded to treatment condition. The same monitoring, video recording, and scoring procedures were applied across the saline- and naloxone-treated groups.

### In vivo assessment of striosome-mediated dopamine suppression during fentanyl withdrawal

To determine whether CRFR1 signaling contributes to withdrawal-induced potentiation of striosome-mediated dopamine suppression, MOR-Cre mice expressing ChRmine in MOR⁺ striosomal neurons and GRAB_DA3h_ in the dorsal striatum were used for simultaneous optogenetic stimulation and fiber photometry recording. On the test day, mice received an intraperitoneal injection of saline or a CRFR1 antagonist. Ten minutes later, dopamine responses to optogenetic activation of MOR⁺ striosomal neurons were recorded for 15 min under baseline conditions. Mice then received fentanyl, and stimulation-evoked dopamine responses were recorded for another 15 min. Thirty minutes after fentanyl injection, naloxone was administered to precipitate withdrawal. Ten minutes after naloxone injection, dopamine responses to the same optogenetic stimulation were recorded for an additional 15 min.

The same stimulation intensity and timing were used across baseline, fentanyl, and naloxone-precipitated withdrawal conditions. Dopamine signals were analyzed as fluorescence changes from baseline, and stimulation-evoked dopamine suppression was quantified during the optical stimulation period. This within-subject design allowed comparison of striosome-mediated dopamine suppression across drug states, while saline- and CRFR1 antagonist-treated groups were compared to determine the contribution of CRFR1 signaling to withdrawal-induced potentiation of dopamine suppression. The fiber photometry experiment was performed using a bundle-imaging fiber photometry setup (Neurophotometrics FP3002 V2). GRAB_DA3h_ sensors were stimulated using 415-nm (isosbestic) and 470-nm excitation LED. The LED power at the ferrule output is between 50 and 100 μW. Z-scores using the following equation:

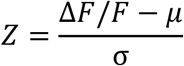

### Histology and cell counting

Animals were anesthetized and perfused intracardially with 4% paraformaldehyde (PFA) in phosphate-buffered saline (PBS). The brains were then extracted and submerged in 4% PFA/PBS solution for one day at 4°C, then transferred to a 30% sucrose solution in PBS. Once the brains had completely sunk in the sucrose solution, they were cut into 50-µm thick coronal sections using a cryostat. The slices were stored in a PBS bath at 4°C before mounting on slides for confocal laser-scanning microscopy (Fluoview, Olympus). c-Fos IHC was carried out on free-floating sections. All incubations (except the primary antibody) and rinses were performed at room temperature on a shaker. Sections were rinsed in PBS three times between steps. Sections were first blocked in 10% bovine serum albumin (BSA) in PBS with 0.3% triton-X (PBST) for 1 hour and incubated in primary antibody (rabbit anti-c-Fos: for mice: 1:2000, EMD-Millipore, #PC38) diluted in blocking solution (1% BSA in 0.3% PBST) overnight at 4 °C. Sections were then incubated in biotinylated donkey anti-rabbit (1:500, Jackson Immuno Research) in blocking solution for 1 hour. Sections were then visualized via incubation for 1 hour in Streptavidin-conjugated Alexa 647 (1:1000, Thermo Fisher) in PBS. All images were processed using Imaris 8.3.1 (Bitplane, Zurich, Switzerland) as previously reported. Immunostaining for DARPP was performed using an anti-DARPP-32 primary antibody, followed by a fluorophore-conjugated secondary antibody emitting at 488 nm, labeling MSNs with green fluorescence. For each brain region, 5-10 brain sections were imaged from each animal. Imaris was used to quantify green and red neurons as well as evaluate colocalization. Brain regions were identified using the Mouse Brain Atlas.

### Slice electrophysiology

Slices were prepared and electrophysiological recordings were conducted as described previously [17, 19, 39, 40, 42–44, 46–53]. Briefly, coronal sections (250 μm) containing the striatum were cut in an ice-cold cutting solution containing (in mM): 40 NaCl, 148.5 sucrose, 4 KCl, 1.25 NaH_2_PO_4_, 25 NaHCO_3_, 0.5 CaCl_2_, 7 MgCl_2_, 10 glucose, 1 sodium ascorbate, 3 sodium pyruvate, and 3 myo-inositol. The solution was saturated with 95% O_2_ and 5% CO_2_. Slices were then incubated in a 1:1 mixture of the cutting and external solutions at 32°C for 45 min. The external solution was composed of the following (in mM): 125 NaCl, 4.5 KCl, 2.5 CaCl_2_, 1.3 MgCl_2_, 1.25 NaH_2_PO_4_, 25 NaHCO_3_, 15 glucose, and 15 sucrose. The external solution was saturated with 95% O_2_ and 5% CO_2_. Slices were then maintained in the external solution at room temperature until use.

Individual slices were transferred to a recording chamber and continuously perfused with the external solution at 2-3 mL/min at 32°C. For CRF-Cre mice infused with ChRmine into the CeA and BNST, red ChRmine fluorescent axonal fibers were visualized using an epifluorescent microscope (Olympus), and striatal neurons were selected within a high-fluorescent area. MSNs were clamped at −70 mV. We used a Cs-based intracellular solution containing (in mM): 125 CsCl, 6 NaCl, 10 HEPES, 1 EGTA, 10 QX-314.Cl, 2 MgATP, 0.6 Na₃GTP, and 2 Na₂CrPO₄. The pH was adjusted to 7.25, and the osmolarity was set at 280 mOsm. To selectively stimulate inputs from channelrhodopsin-expressing fibers onto DMS neurons, 590-nm light was delivered through the objective lens for 2 ms every 20 sec throughout the experiment. After a stable baseline, TTX was bath-applied for 5 min, followed by TTX + 4AP bath application.

For MOR-Cre;Ai14 mice, red striatal neurons were selected. To measure sEPSC, we used a Cs-based intracellular solution containing (in mM): 119 CsMeSO_4_, 8 tetraethylammonium chloride, 15 4-(2-hydroxyethyl) piperazine-1-ethanesulfonic acid (HEPES), 0.6 ethylene glycol tetraacetic acid (EGTA), 0.3 Na_3_GTP, 4 MgATP, 5 QX-314, and 7 phosphocreatine, with an osmolarity of ∼280 mOsm/L. The pH was adjusted to 7.3 with CsOH. The sEPSC was first measured for 3-6 min. We then bath-applied either CRF or a CRF+CRFR1 antagonist for 5 min, then measured sEPSC again for 6 min. All data were analyzed using Clampfit 10.5.

### Statistical analysis

All data are presented as mean ± standard error of the mean (SEM). Normality was assessed using the Shapiro-Wilk test. If the data met normality assumptions, statistical comparisons were conducted using a two-tailed t-test (paired or unpaired) or one-way ANOVA (with repeated measures), followed by post hoc Tukey’s or Sidak’s multiple comparisons test, when appropriate. If normality was not met, data were analyzed using the Mann-Whitney test or Kruskal-Wallis test (followed by Dunn’s post hoc test). Statistical significance was set at p < 0.05. All statistical analyses were performed using SigmaPlot, and graphs were generated using OriginPro.

## ACKNOWLEDGEMENTS

We appreciate critical comments on our manuscript from Wang lab members. This research was supported by NIAAA Grant R01AA021505, R01AA027768, U01AA025932 and X-grant from Texas A&M University to J.W., McGovern Fellowship from Texas Research Society on Alcoholism (TRSA) to X.X., and Doctoral Student Small Grant from Research Society on Alcohol (RSA) to X.X.

## Author contributions

Conceptualization: J.W., X.X.; Methodology: X.X., J.W.; Investigation and Formal analysis: X.X., A.E., R.C.; Writing - Original Draft: X.X.; Writing – Review & Editing: J.W., X.X.; Resources: X.W.; Funding acquisition: J.W. and X.X.; Supervision: J.W.

## Declaration of interests

The authors declare no competing interests.

**Declaration of generative AI and AI-assisted technologies in the writing process** During the preparation of this work, the authors used ChatGPT (OpenAI) to improve the grammar and readability of the manuscript. After using this tool, the authors reviewed and edited the content as needed and took full responsibility for the published article.

## Notes

### Competing Interest Statement

The authors have declared no competing interest.

